# Comparing host module activation patterns and temporal dynamics in infection by influenza H1N1 viruses

**DOI:** 10.1101/2021.05.07.443162

**Authors:** Irina Nudelman, Daniil Kudrin, German Nudelman, Raamesh Deshpande, Boris M. Hartmann, Steven H. Kleinstein, Chad L. Myers, Stuart C. Sealfon, Elena Zaslavsky

## Abstract

Influenza is a serious global health threat that shows varying pathogenicity among different virus strains. Understanding similarities and differences among activated functional pathways in the host responses can help elucidate therapeutic targets responsible for pathogenesis. To compare the types and timing of functional modules activated in host cells by four influenza viruses of varying pathogenicity, we developed a new DYNAmic MOdule (DYNAMO) method that addresses the need to compare functional module utilization over time. This integrative approach overlays whole genome time series expression data onto an immune-specific functional network, and extracts conserved modules exhibiting either different temporal patterns or overall transcriptional activity. We identified a common core response to influenza virus infection that is temporally shifted for different viruses. We also identified differentially regulated functional modules that reveal unique elements of responses to different virus strains. Our work highlights the usefulness of combining time series gene expression data with a functional interaction map to capture temporal dynamics of the same cellular pathways under different conditions. Our results help elucidate conservation of the immune response both globally and at a granular level, and provide mechanistic insight into the differences in the host response to infection by influenza strains of varying pathogenicity.

## 1 INTRODUCTION

The possibility of influenza virus pandemics remains a potent public health threat. While most annual influenza strains are associated with a relatively low global infection rate and mortality, more widely infectious or lethal influenza virus strains arise periodically. The influenza pandemic of 1918 was responsible for more than 50 million deaths and, within one year, reduced the life expectancy in the United States by a dozen years (1). More recently, the swine-origin influenza pandemic in 2009 infected 20-50 percent of the population of some countries, although, fortunately, it had a mortality rate comparable to that of seasonal influenza strains (2). Thus, individual seasonal and pandemic influenza strains vary in their infectivity and pathogenicity. Although the genetic mechanisms underlying the emergence of new viruses are relatively well understood, less is known about virus-host interaction effects that may influence influenza transmission or disease outcome. Implementing a computational approach to identify commonalities and differences in the host biological response to different influenza virus strains is important in providing insight into common and distinct components of the host response program that may contribute to pathogenicity.

Increasingly, emerging research suggests that temporal dynamics may play an important role in the varying pathogenicity that is observed among different influenza strains (3). This premise motivates a systematic study of time series expression datasets to gain a more complete understanding of the differences in host response dynamics observed with each virus. However, time series analyses present computational and experimental challenges. Measurements must be obtained at the appropriate time scales. Proper temporal alignment among different datasets and possible time shifts in activity patterns need to be addressed when interpreting such data. The standard approach of identifying lists of differentially expressed genes provides only limited insight into the biological mechanisms underlying commonalities and differences among host responses to multiple influenza strains (4, 5, 6, 7, 8, 9).

Integration of gene expression data with complementary information about physical or functional associations between molecular entities has been proposed as a powerful approach to improve the interpretation of global transcriptional changes. These integrative approaches analyze gene expression experiments in the context of an independently constructed connectivity map, such as a protein-protein interaction (PPI) network, to identify modules comprised of genes or proteins that participate in common biological pathways or functions (10). More recently, integrative methods have been developed to identify ‘active’ modules (i.e. related groups of genes exhibiting concordant transcriptional changes (10, 11, 12, 13)), modules conserved across species (14, 15) and ‘differential’ modules (16, 17). (For an overview, see also a review article by T. Ideker and colleagues (18) and references therein).

Time-course gene expression datasets capture important features of the temporal trajectories of transcriptional changes. While the majority of integrative gene expression and interaction network analyses have not utilized the temporal dimension of the data, there have been attempts to incorporate temporal information into module discovery (19, 20, 21, 22, 23, 24). For example, Gao and Wang (22) used a phase-locking approach (25) to identify yeast cell cycle genes that show temporal coordination and whose interactions are supported by a PPI network. In another study, Jin and colleagues (23) applied a time-warping dynamic programming algorithm (26) to identify locally-similar temporal expression patterns among groups of genes forming connected components of a PPI network. These methodological advancements do not offer a solution to the problem we call *‘comparative module discovery’*, i.e. the identification of temporally-shifted, network-based patterns of expression showing conservation (or divergence) *between* time-course datasets that are generated in the same experimental system by different perturbations. Developing such an analysis method would be valuable in elucidating commonalities and differences in the biological responses to these perturbations. The identification of such comparative modules is critical for addressing the central question of our study - that of understanding the similarities and differences in virus-host interaction effects in response to related influenza virus infections.

In order to perform comparative module discovery, we developed a novel integrative DYNAmic MOdule (DYNAMO) method, and applied it to understand the common and unique features of the host immune response to infection by related strains of the influenza virus. Integrating datasets that capture the temporal progression of the global gene expression response post-infection with an interaction network, our method discovers both conserved and differential comparative modules. Conserved comparative module discovery identifies a set of highly functionally connected genes that show a high degree of similarity between their regulation and response patterns for perturbations being compared. Our approach allows the possibility that the module responses may be shifted in time across different perturbations. Differential comparative module discovery identifies genes that show differences in their pattern of regulation across different perturbations. Differential module discovery is a difficult problem because truly condition-specific regulatory patterns must be distinguished from experimental and biological variability (18, 14). Our method is able to identify high-confidence differential subnetworks by exploiting the temporal nature of the expression data and anchoring the modules in functional network connectivity relationships. By computing the optimal temporal alignment of each module’s expression profile between two different conditions, we are able to capture divergent activation patterns. Overall, our method addresses the broad problem of combining functional connectivity and genome-scale time series expression data to extract vital temporal information and to enable a comparison of gene programs and module activation across time.

We apply DYNAMO to the problem of studying host-pathogen interactions for multiple H1N1 influenza virus strains. Our study builds upon the availability of identically sampled time series data for H1N1 seasonal and pandemic influenza virus of a human immune cell that lends itself to a systems-wide comparison of the dynamics underlying the modulation of the host response by each virus (27). DYNAMO extracts functionally conserved modules that show a difference in their temporal dynamics or pattern of transcriptional changes between each pair of infection time-course datasets. We demonstrate that the groups of modules identified are statistically significant and that the algorithmic element of optimization for the best temporal alignment is crucial for their identification. Our analysis provides insight into the biological mechanisms underlying the module response patterns elicited by these influenza virus strains. DYNAMO is accessible via a user-friendly interface at http://tsb04.mssm.edu/.

## 2 MATERIALS AND METHODS

### 2.1 Subnetworks with shifted temporal dynamics

DYNAMO searches for groups of genes in two time-series expression experiments that exhibit similar gene-by-gene expression patterns while allowing a temporal shift. DYNAMO is an integrative method that overlays expression data on a functional interaction network and leverages the methodology of the neXus algorithm (14) to reinforce functional coherence within each discovered module.

#### 2.1.1 Overview of the neXus algorithm

The neXus algorithm (14) was developed to search for conserved subnetworks between a pair of expression datasets across species or within a single species. In its single species version, the method attempts to form dense gene subnetworks within a chosen functional interaction network while maintaining sufficient similarity in the expression levels of the subnetwork genes. Briefly, neXus initiates a depth-first search from a seed gene as it aims to build a subnetwork in the underlying functional network. For each gene that it considers for addition to the growing subnetwork, two conditions have to be met. First, it evaluates that the connectivity requirement is met by maintaining a minimum desired clustering coefficient of the genes in the putative subnetwork. Second, the expression similarity condition is evaluated by computing the average expression activity score of the subnetwork genes. The subnetwork grows until there are either no genes to be considered or it has reached a maximal size. The process is repeated for every initial seed gene, and final subnetworks with large overlaps can be merged.

#### 2.1.2 Incorporating a time shift

Consider the expression vectors of gene *g* in two aligned time-course expression datasets. DYNAMO evaluates optimal similarity between the two vectors while allowing one vector to be shifted relative to the other by some time shift, Δ*t*. To assess similarity in expression at any such Δ*t*, we calculate time-lagged Pearson correlation coefficient of the two vectors. Let *T* be the set of discrete time points at which gene expression was sampled for each virus infection and *T* ^*′*^ be the corresponding set of time points shifted by Δ*t*. Denoting the expression vectors as *XT* (*g*) for the stationary time-course and *Y*_*T*′_ (*g*) for the time-shifted course, we compute time-lagged correlation coefficient, (TLC) 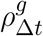 for gene *g* between the two responses as

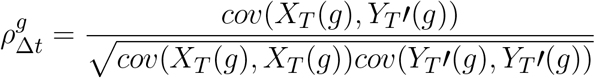

where *cov* is the standard covariance. We use linear interpolation to calculate the values in the stationary time course that correspond to the new time points. Just like the standard correlation, a time-lagged correlation close to 1 means that the expression of gene *g* is perfectly correlated between the two responses once the time-shift is taken into account. We determined (data not shown) that transforming the correlation distributions via the Fisher Z-transform

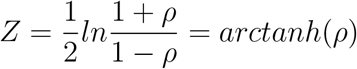

resulted in better findings, and used these Fisher-transformed scores within the algorithm when assessing expression coherence of growing subnetworks at various time lags.

#### 2.1.3 Algorithm to find temporally-shifted subnetworks

We begin with a list of seed genes, and their expression vectors from a pair of aligned time-course experiments. We use fold-change values over a control condition, though other quantitative vectors such as differential expression p-values can be used as well. The matrix of standardized z-scores is computed for all genes at every considered time lag. Putative subnetworks are grown greedily from every seed in turn. First, candidate genes are identified via a depth-first search from the seed gene, as in the original neXus algorithm (14). To assess subnetwork coherence at a particular time lag Δ*t*, we calculate the *subnetwork’s score* as the average of the Fisher-transformed z-scores of its constituent genes at that time lag. To select a gene for addition to the growing subnetwork, DYNAMO then optimizes over all considered time lags and corresponding putative subnetwork scores, provided that the minimum desired connectivity requirement in the underlying functional network is met. The connectivity condition, measured as the average weighted clustering coefficient of the subnetwork, enforces functional coherence of the growing module. The network score maximization component enables the algorithm to identify the best time-lag (if one exists) that brings the group of genes in the two responses into temporal alignment. Note that the optimal time lag for a growing network can change with addition of new genes, but, in our experience, does not vary widely. We use the average of the per-gene maximal fold-changes during the time course in each response as a third cutoff to be met in order to filter out false high subnetwork scores that may be due to a good alignment of flat time courses of genes that do not show significant differential expression. Finally, we merge the discovered subnetworks if there is considerable (0.6) overlap among their constituent genes and their identified time lags are the same.

### 2.2 Subnetworks with differential expression patterns

Identifying genes that behave differently between a pair of responses is a difficult problem because many spurious expression differences can arise for individual genes. We again employ the insight of constraining expression differences by requiring tight clustering of such genes in the underlying functional network. The structure of the algorithm is similar to that of the algorithm for finding conserved temporally-shifted subnetworks. We enforce the network connectivity requirement by maintaining a minimum desired clustering coefficient, and optimize the choice of candidate genes for addition to the growing subnetwork by selecting one that shows the highest divergence in its expression pattern between the responses, provided that the average expression score stays below a selected score threshold. The subnetwork expression score that, in the case of differential modules, needs to identify genes with divergent expression patterns, is modified to reflect that difference. We observe that the correlations and their corresponding Fisher z-score distributions for most time lags have positive means (Supplementary Figure 1), indicating that most genes show similar expression trends in the infection responses. Genes that exhibit different expression trends between responses may show only slightly negative absolute correlations. To better identify such genes, we use standardized rather than raw Fisher z-scores. This way, DYNAMO searches for genes that are within some number of standard deviations below the mean of the Fisher z-score distributions. The average fold-change requirement is also altered to enforce that only subnetworks in one of the responses pass the cutoff. This change allows the algorithm to capture both subnetworks that show opposing activation patterns as well as those that show activation in only one of the responses.

### 2.3 Assessing subnetwork significance

We employ a randomization analysis, and use it as a tool to calibrate various DYNAMO parameters and assess biological significance of the subnetworks discovered in the comparison of the influenza infection responses. We create five randomized expression profiles by randomly shuffling the expression vectors with respect to gene labels. Our algorithm for subnetwork discovery is applied to these profiles while the functional network structure remains intact, enabling an estimation of the temporally shifted (or divergent) expression pattern coherence that arises from the clustering of genes by random chance. For a given expression score threshold, the subnetworks discovered in the randomized data at that threshold represent false positive findings and enable an estimation of false discovery rate. We calculate the associated subnetwork confidence value as

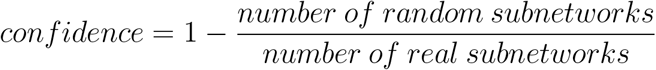

and use it to assess the subnetworks’ statistical significance. Overall, exploring the algorithm’s findings over various parameter ranges for randomized and real data allows a substantiation of our parameter choices and a quantification of the biological significance of the results.

### 2.4 Experimental data and algorithm parameters

#### 2.4.1 Microarray data

Human monocyte-derived dendritic cells were infected with each of the four strains of the H1N1 influenza virus (Tx, NC, Brevig and Cal). For each infection, cells were collected at the following time points post infection: 120, 160, 200, 240, 300, 360, 420, 480 min. Näive non-infected DCs underwent the same experimental handling as infected DCs in virus-free allantoic fluid to ensure that mechanical manipulations could not be responsible for differences in experimental readouts. These served as a negative control time-course. All time points and controls were performed in triplicates. The details of DC maturation, virus preparation and infection as well as RNA extraction for microarray experiments are described elsewhere (27). The RNA samples were processed and hybridized to HumanHT-12 v4 Expression BeadChip Kit (Illumina San Diego, CA) by the Yale Center for Genome Analysis following the manufacturer’s instructions, and raw expression data were output by the Illumina GenomeStudio software. These data were log-transformed, filtered for minimum intensity (*log*_2_(*expression*) *>* 6.6), determined based on visual inspection of the distribution), averaged over the triplicates, and converted to fold-change values over the time-matched allantoic fluid control condition. Each viral time-course was analyzed for differential expression using LIMMA (BioConductor implementation) after correction for multiple hypothesis testing (*q <* 0.05) (28). Maximally expressed probes were chosen for differentially expressed genes with multiple probes. We took the union of genes that passed the differential expression criterion at any time-point in each viral time-course as our candidate seed set.

#### 2.4.2 Flow Cytometry

Human monocyte-derived DCs were infected with either NC, Tx (both seasonal) or Cal (pandemic) H1N1 IAV. Samples were fixed in 1.6% paraformaldehyde (Sigma) and subsequently stained with fluorophore conjugated antibodies against CD86 and HLADR (both BD) at multiple time points post infection. Cells were analyzed with a LSRII flow cytometer (BD) and data was analyzed with Cytobank and R.

#### 2.4.3 Functional networks

We consider two human functional linkage networks, the general network (29) that is trained on diverse curated functional pathway data (30) and an immune-specific network (31) trained on immune pathways only. Both networks integrate many heterogeneous data from diverse sources including physical and genetic interactions as well as microarray data to create the functional associations between gene pairs. The edge weight distributions, which reflect the confidence in the gene-gene associations and are inferred via Bayesian integration, differ between the two networks, with the median edge weights being 0.85 and 0.22 for the general functional network and the immune-specific network, respectively. We retained one million most highly weighted edges for each network. We explored the algorithm’s performance and its dependence on the clustering coefficient parameter for each network separately (data not shown), and found that in each case there exists a range of this parameter (different for each network because of the differences in the underlying edge weight distributions) with comparably good performance. We use these ranges, and set the average clustering coefficient cutoffs to 0.8 and 0.5 for general and immune-specific networks respectively.

#### 2.4.4 Algorithm parameters

We chose the values of 1.5 for subnetwork score (see discussion in Supplementary Materials), 0.5 for minimum clustering coefficient of the immune-specific network and 0.9 for subnetwork confidence, and considered subnetworks that pass all these cutoffs. The subnetworks were grown to maximal size of 25 nodes. Additional internal neXus parameters were left at their defaults.

## 3 RESULTS

### 3.1 Overview of algorithm developed for comparative module discovery

We developed DYNAMO to find conserved and differential subnetworks that exist between time-course datasets that measure gene expression responses to different perturbations in the same experimental system. As a motivation for our study was an observation made while investigating a time-course microarray dataset of the responses to four strains of the influenza virus in human monocyte-derived dendritic cells (DCs) (27). We noted a time shift in the expression dynamics for many important immune response genes (Figure 1). This served as an impetus for addressing the aspect of temporal dynamics in our methodology development. The DC infection study comprised of four A/H1N1 influenza viruses that differ in their infectivity and clinical severity, including two pandemic strains, the influenza of 1918 (Brevig) and the recent 2009 strain (Cal), as well as two seasonal strains, the New Caledonia strain of 1999 (NC) and Texas 1991 (Tx). Global expression was sampled with high frequency during the first eight hours post-infection. Thus, our investigation relied on a well-controlled dataset representing time-course responses in a single cell type to antigenically similar influenza strains varying in pathogenicity and transmission efficiency.

**Figure 1.**
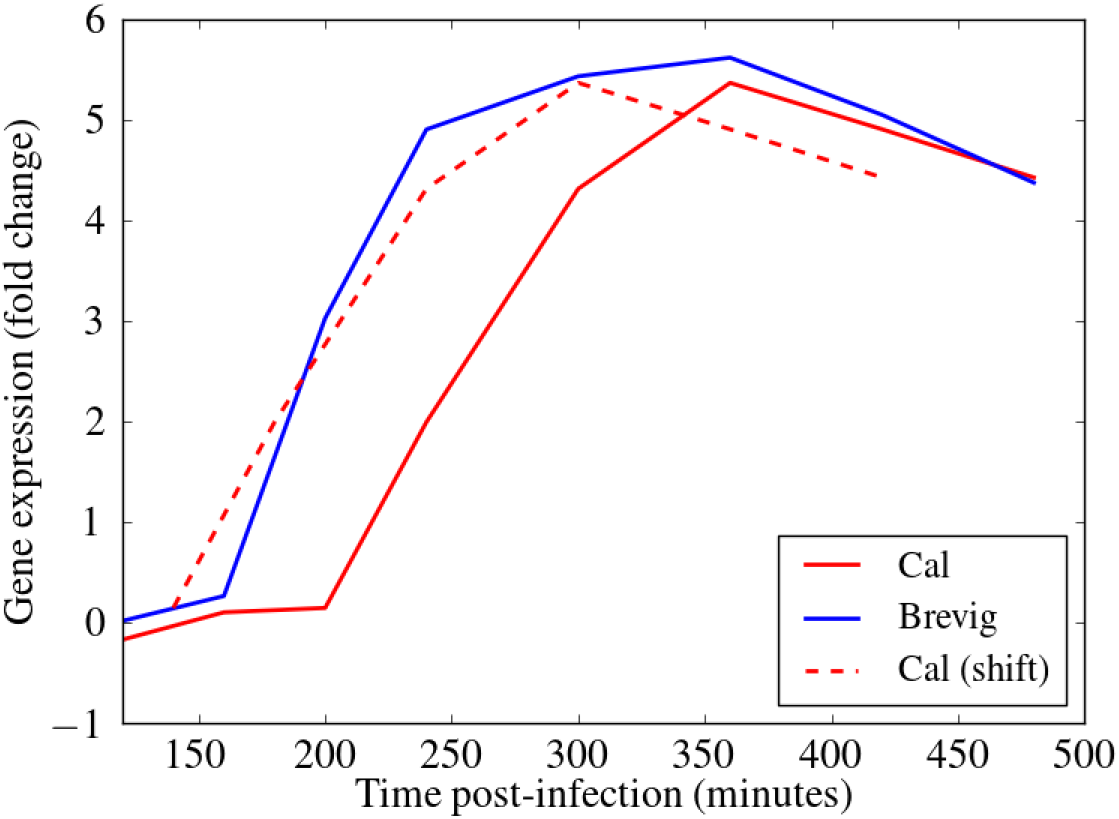
Alignment of the gene expression time courses for MX1, an important immune response gene (51), following Cal and Brevig Influenza H1N1 infections. The solid red and blue lines represent the actual MX1 expression in Cal and Brevig responses, respectively. The dashed red line represents the Cal time course shifted to the left by 60 minutes. The fold change gene expression measurements are normalized relative to the results obtained from uninfected control cells.

We built upon the approach of neXus (14), an algorithm that overlays gene expression data on functional interaction networks (29, 32) to identify functionally coherent groups of genes that have similarity in their expression patterns across species. We describe our approach briefly here, and a more extensive discussion of the algorithm is found in the Materials and Methods section. Each DYNAMO module is grown from a seed gene by adding nearby genes in the interaction network in a way that maximizes the average gene expression activity score of the module, while maintaining a minimum desired clustering coefficient. DYNAMO’s expression activity score (subnetwork score) addresses the challenge of comparing time-course datasets and studying response programs that may be temporally shifted with respect to one another. DYNAMO samples time-shifts in the gene expression dynamics, computing the time-lagged Pearson correlation coefficient, and conducts a greedy search for coherent active subnetworks, such that each module member gene in one dataset exhibits a maximally similar expression pattern (possibly with a temporal shift) to the same gene in the other dataset. For each module, the optimal time shift, applied to all genes, is identified. Subnetworks with high overlap in gene membership that exhibit the same time lag are merged. DYNAMO identifies the set of highly coherent, statistically significant modules by determining the false discovery rate (FDR) via analysis of randomly shuffled expression data. The same methodological approach is applied to the problem of differential comparative module discovery. DYNAMO identifies maximally differentially regulated genes in two datasets that represent a highly functionally related module in the underlying functional interaction network.

In the following sections, we used DYNAMO to identify and compare modules in time-course responses to the different influenza viruses. We first performed an in-depth analysis of the Brevig/Cal response comparison, validating our method and offering insight into the biology of their shared and unique response processes. We then compared temporal dynamics and functional pathway activity, computed as GO term enrichment of discovered modules, for all the strains. Detailed analyses of each comparison, including conserved and differential comparative modules, functional pathway activity and performance characteristics are available at http://tsb04.mssm.edu/.

### 3.2 Evaluating the DYNAMO algorithm

We evaluated two important aspects of the DYNAMO algorithm. First, we considered the effect of the choice of the functional network used by DYNAMO to identify functional connectivity. Next, we assessed the effects of allowing a temporal shift of the gene expression dynamics on module discovery. In evaluating the algorithm’s performance, we considered the number of conserved modules that were discovered by the algorithm, and we estimated the false positive rates for the discovered modules via a randomization analysis (see details in the Materials and Methods).

#### 3.2.1 Functional network selection

Functional networks are constructed from heterogeneous data sources and represent diverse associations between genes or proteins (33, 29). Bayesian integration of multiple data types, including protein-protein and genetic interactions, gene expression, protein localization, phenotype, and sequence data, was used to infer functional connections between molecular entities. Given their more comprehensive coverage of a broad variety of gene relationships, functional networks allow for more sensitive discovery of conserved active modules and have been shown as advantageous for this task over protein-protein interaction networks (14).

We assessed DYNAMO’s performance using the general human functional network (29) and an immune-specific functional network that should, in principle, capture associations that are more relevant for immune contexts (31). The edge weight distributions, which reflect the confidence in the gene-gene associations and are inferred via Bayesian integration, vastly differ between the two networks. Correspondingly, the network related parameters can not be set at the same values for the two networks. Retaining one million most confident edges for each network, we explored the algorithm’s performance for each network separately, and set the clustering coefficient parameters to values that maximized performance for each network individually (see Materials and Methods for details). We identified the conserved comparative modules for the Brevig and Cal pair using the two functional networks and varying the gene expression based activity score parameter of the algorithm. To assess the statistical significance of the modules discovered, we performed a randomization analysis. Specifically, the expression time-course vectors were randomly shuffled five times with respect to the gene labels, and the algorithm was applied to the shuffled expression profiles. Any modules identified in these randomized expression data would represent false positives and not biologically meaningful conservation. As seen in Figure 2, utilizing the immune-specific functional network was far superior to using the general functional network. We observed that many more coherent modules were discovered at every subnetwork score cutoff, suggesting that the functional connectivity that underlies the gene relationships in influenza responses is better reflected in the immune-specific functional network. Focusing on the randomization experiments, our evaluations suggested a false discovery rate of *<* 5% for a broad range of subnetwork score cutoffs when using the immune-specific functional network. Overall, comparing DYNAMO’s results with the two different underlying functional networks, we found that many more modules are discovered at every FDR setting when the immune-specific network is used, indicating in a substantial improvement in sensitivity.

**Figure 2.**
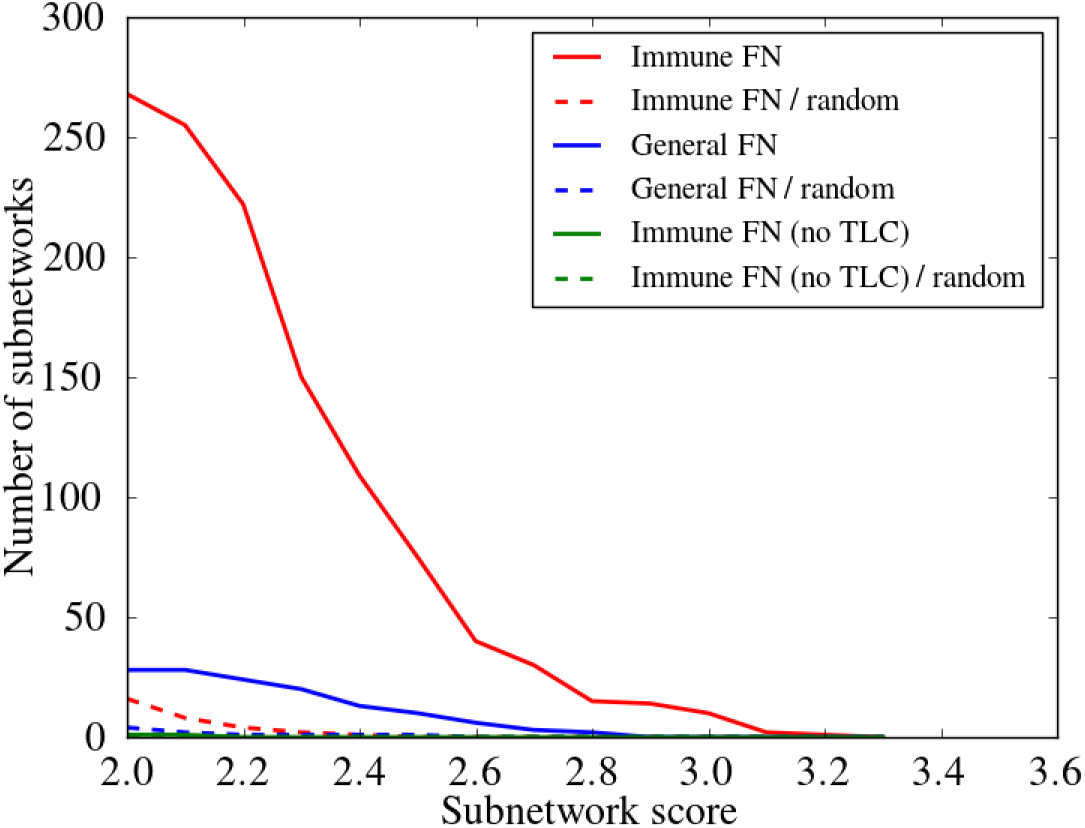
Assessment of the choice of the functional network (FN) and importance of introducing a temporal shift in conserved comparative module discovery. The number of subnetworks across a wide range of subnetwork expression scores (see Materials and Methods), is compared for different approaches. To enforce functional coherence, two different functional networks were used, a general (General FN) and an immune-specific (Immune FN) functional network. Subnetwork identification was performed employing either standard (no time lag) or optimized time-lagged Pearson correlation (TLC). A randomization analysis, averaged over five randomization instances, was performed for each comparison to assess the false discovery rate (dashed lines). The red curves are produced using TLC optimization and Immune FN, the blue curves use TLC optimization and General FN, and the green curves use standard Pearson correlation and Immune FN. All discovered modules are reported, regardless of the confidence threshold. Note that the number of modules discovered never rises above one for the green curves.

We further assessed the importance of enforcing the functional coherence of the modules and considered whether our method can extract high-confidence subnetworks from expression data alone. We used DYNAMO without enforcing the clustering coefficient parameter, while adding putative module member genes in the same order from a pool that is functionally proximal to the seed gene. As shown previously by Deshpande et al. (14) and corroborated in our analysis, fewer modules were discovered without enforcing the clustering coefficient parameter. Furthermore, they were of low significance in view of the similar number of modules identified via a randomization analysis. We concluded that using a functional connectivity map and selecting a map that is most relevant for the experimental study (an immune-specific functional network in our case of subnetwork discovery in the responses to influenza viral infection) are essential for identifying significant modules. As such, we used the immune-specific functional network in all further evaluations within this study.

#### 3.2.2 Advantage of allowing a temporal shift

We evaluated the advantage gained by the introduction of a time shift in the identification of active subnetworks shared by the Brevig and Cal responses. We considered possible time lags of -80, -60, -40, -20, 0, 20, 40, 60 and 80 minutes, and shifted the Cal time-course with respect to the Brevig time-course. We compared DYNAMO’s results when optimizing module discovery over the possible time-lags to those found with no time shift allowed (i.e. using standard Pearson correlation), while keeping all other parameters the same. As shown in Figure 2 and noted above, many conserved temporally shifted subnetworks were identified over the range of considered network score cutoffs (red curve). In contrast, almost no subnetworks were identified when a temporal shift was disallowed (green curve). These observations indicate the importance of the temporal shift element in enabling discovery of conserved comparative functional modules.

### 3.3 Comparison to existing algorithms

DYNAMO’s objective in identifying conserved or divergent temporally shifted modules that are common between two responses is quite unique, and, to the best of our knowledge, has not been addressed in the literature. Nonetheless, we evaluated DYNAMO against two other methods that are most similar and identify conserved subnetworks from gene expression data, ModuleBlast (34) and TDARACNE (35).

ModuleBlast was designed to compare module activation patterns across species. It uses expression data and network topology information to search for conserved and divergent sub-networks. Analysis of the host immune response gene expression data comparing Brevig and Cal infections using ModuleBlast resulted in 38 modules. These modules were generally not functionally enriched for immune-specific processes, according to functional annotation within ModuleBlast. Analysis by DYNAMO shows the importance of the network context in which gene expression data is analyzed. Biological pathways that are activated in an immune context are best identified using an underlying network that emphasizes immune-specific interactions. Since ModuleBlast employs a generic interaction network, the relative paucity of conserved modules is not surprising. Furthermore, while ModuleBlast makes use of temporal information, it does not optimally align the responses. This is a key difference that enables DYNAMO to capture coherent activation patterns that are temporally shifted.

We also applied TDARACNE to our dataset. TDARACE was designed to address a different problem - it is a subnetwork inference method that is not comparative and operates on each gene expression dataset individually. Therefore, it generally infers dissimilar sets of modules for the Brevig and Cal datasets, making a direct comparison with DYNAMO meaningless.

### 3.4 Conserved time-shifted host response to Brevig and Cal influenza strains

DYNAMO identified 207 high confidence functionally-coherent subnetworks that are time-shifted between the two pandemic strains, Brevig and Cal. To evaluate the subnetworks for biological significance, we assessed functional enrichment in the set of genes contained in each subnetwork. The enrichment was computed for each subnetwork individually based on the overlap of its constituent genes with the Gene Ontology (GO) (36) biological process terms using enrichR (37). Every subnetwork identified was enriched for at least one GO term with p-value of 0.0001. Overall, 71 GO terms were associated with the discovered subnetworks, of which vast majority were immune related. Nearly all subnetworks were annotated with GO terms describing cytokines, type I interferon signaling, and response to virus (Figure 3). This enrichment is consonant with extensive experimental evidence identifying cytokine and interferon responses elicited by influenza infection (38, 39, 6, 8). In particular, type I interferons provide a first line of defense against the virus (Garcia-Sastre et al., 1998), functioning in both autocrine and paracrine fashions to prevent its replication and spread to neighboring cells, and are crucial in host defence against influenza infection.

**Figure 3.**
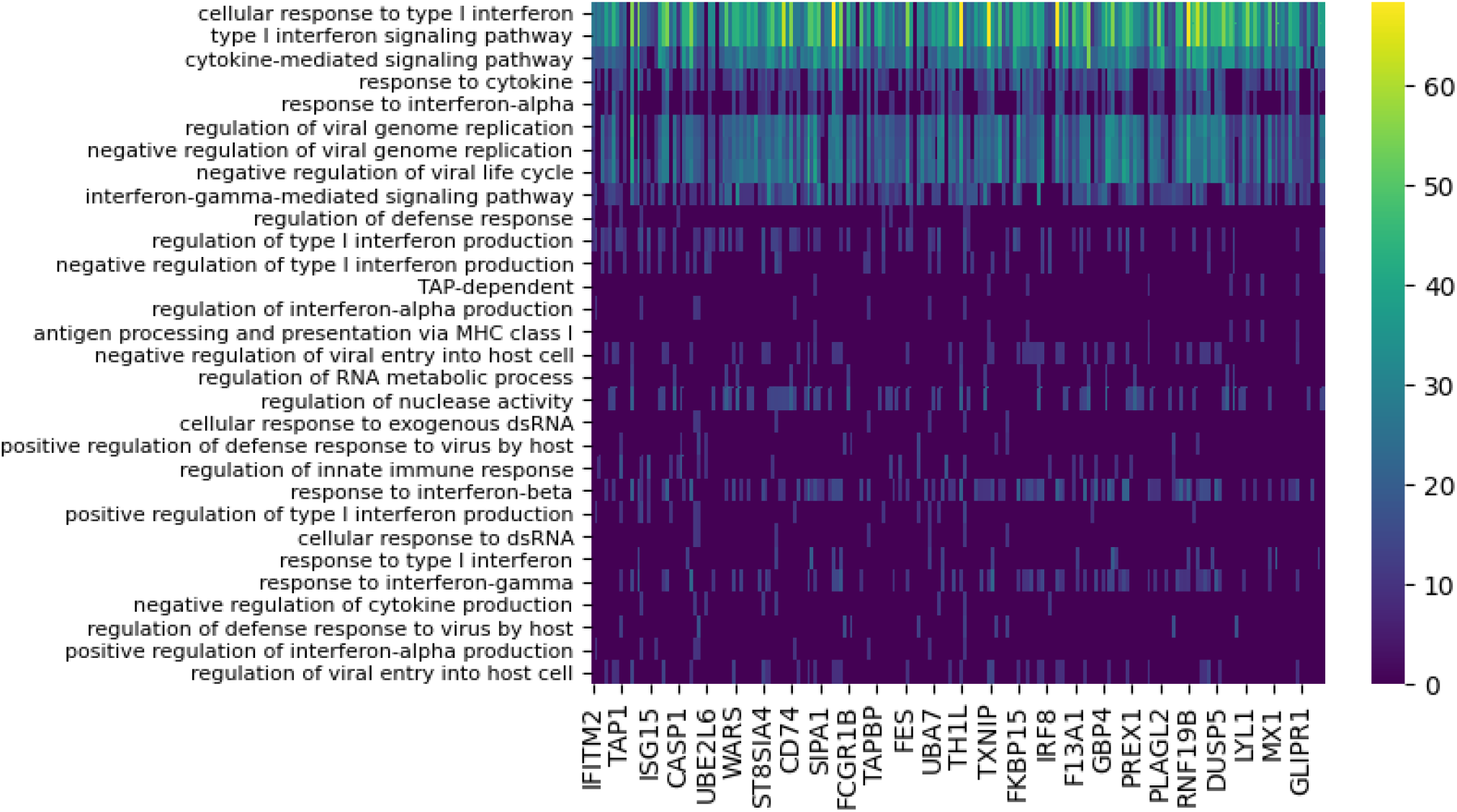
Functional enrichment analysis of conserved subnetworks appearing with the 80 minute time lag for the Cal/Brevig comparison. The immune-specific nature of the modules was determined by using GO term enrichment for individual subnetworks with p-value cutoff 0.0001. The rows represent the GO terms and the columns are the individual subnetworks, indexed by their seed genes. The color scale indicates the level of significance of the GO term enrichment and is based on -log(pvalue). For clarity, GO terms that assign to fewer than seven subnetworks or that annotate more that 100 genes in GO are not shown.

Absolute majority of the subnetworks (82%) identified showed optimal similarity when aligned at the 80 minute time lag, with the Cal response activated after the Brevig response. Our findings confirmed the earlier observations (3) that the highly pathogenic Brevig strain is characterized by rapid activation of the host immune response and that this early activation may account for the extreme severity of disease caused by this strain (40). Furthermore, the considerable similarity in the activated immune response program when accounted for the shift in temporal dynamics indicated that the timing of the host immune response may be at the basis of the key differences observed between disease outcomes for these two infections.

### 3.5 Conservation and temporal alignment of the global host response and specific immune processes

We next used DYNAMO to identify conserved temporally shifted modules to compare all pairs of influenza strains. Table 1 summarizes the results for each pair-wise comparison and includes the dominant time lag, i.e. the time lag assigned to the largest fraction of the discovered subnetworks. For example, for the Brevig/Cal pair, the Brevig response is shifted 80 minutes earlier in comparison to Cal for the majority of modules identified. We found that the responses to the two seasonal strains, Tx and NC, show the largest number of subnetworks, and 69% of them show maximal similarity with no time shift. The increased number of similarly regulated subnetworks in the Tx/NC comparison resulted from a large down-regulation effect not seen with the other viruses (27). Because the algorithm optimizes subnetwork conservation over time and different components of the responses may contribute to conservation for each strain pair, some dominant time lags appear inconsistent. For example, the Tx/NC pair exhibits the dominant time lag of zero, but these responses do not have the same time lag relative to the Cal strain. However, a clear overall temporal pattern emerges. The conserved modules show that the Brevig infection elicits the earliest response, Tx and NC are intermediate, and Cal is the latest.

**Table 1.** Subnetworks conserved across pairs of influenza responses. For every comparison, the lagging influenza strain response is listed second (e.g. the Cal response is delayed compared to Brevig). The dominant time lag is measured in minutes, and the last column indicates the percent of modules that are found at the dominant time lag.

| Virus comparison | # subnetworks | average size | dominant time lag | % at dominant time lag |
| --- | --- | --- | --- | --- |
| Brevig/Cal | 207 | 24 | 80 | 83 |
| Tx/Cal | 377 | 21 | 20 | 54 |
| Tx/NC | 2982 | 24 | 0 | 69 |
| Brevig/Tx | 382 | 22 | 20 | 60 |
| NC/Cal | 203 | 24 | 80 | 83 |
| Brevig/NC | 400 | 22 | 20 | 37 |

An overall conservation of the immune response for all the pairwise comparisons was evident in the functional enrichment observed in the subnetworks. Using GO term enrichment by enrichR (37), we found a set of 27 highly enriched (*p <* 0.0001) immune-related GO terms that were common to all the comparisons and collectively were assigned to the absolute majority of the subnetworks. Representative GO terms are shown in Figure 4. They describe host immune response to viral infection and capture cytokine and interferon-regulated processes that are essential in defense against the influenza virus. For each comparison individually, these processes exhibited a temporal consistency, assigning to modules with a singular time lag. These results imply that there is a conserved set of immune processes that is activated in response to the four different influenza strain infections. Within the response to one infection, the relative timing of these immune processes appears to be consistent. In comparisons of the responses, these coherent immune processes are shifted in time on block (Figure 4). This suggests the presence of a highly conserved core of the host immune response.

**Figure 4.**
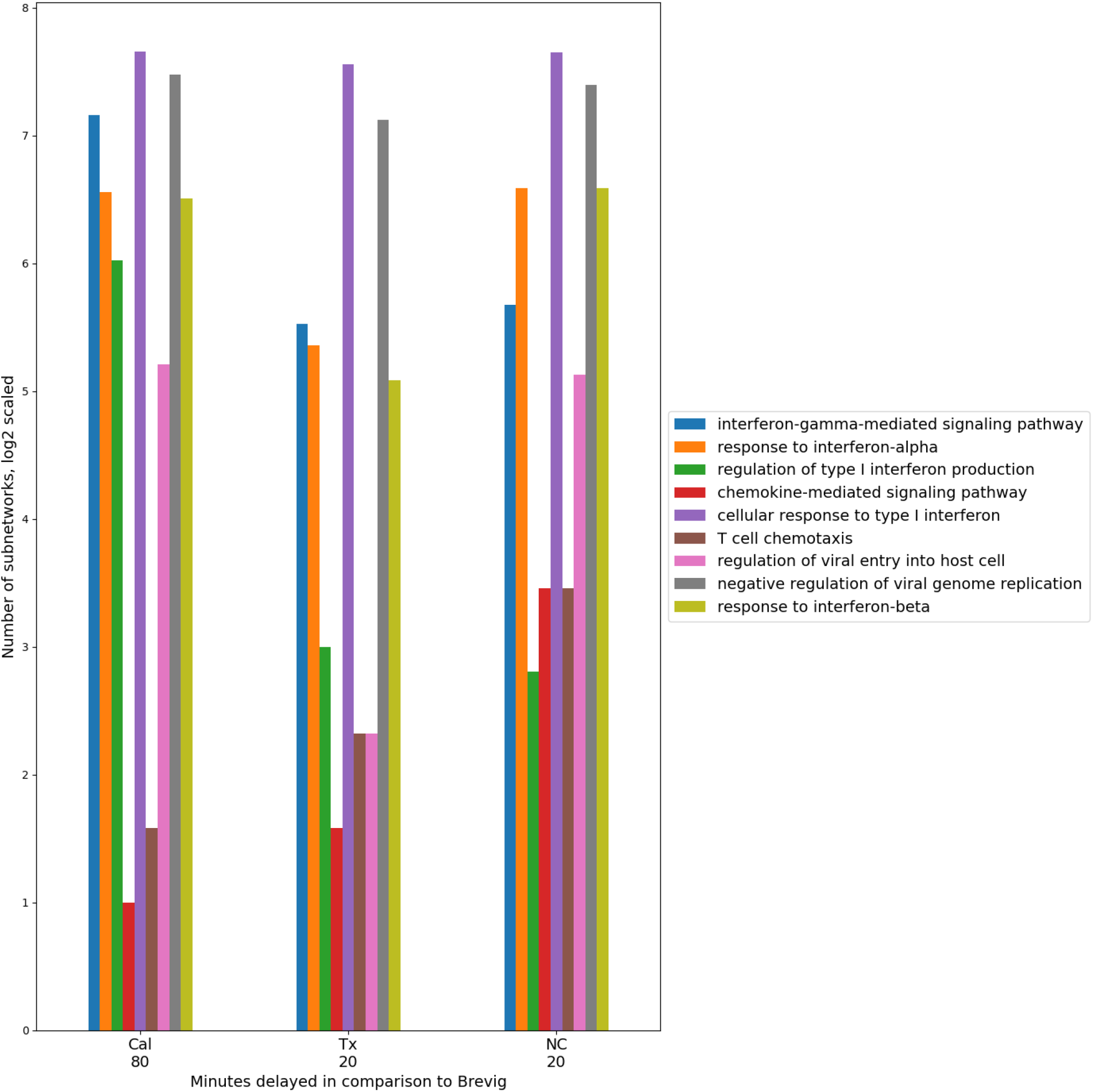
Timing consistency of the nine conserved immune response processes in the pair-wise comparisons between Brevig and the other virus strain responses. A group of representative GO terms was selected from the larger immune-related set of 27 processes found enriched among conserved modules for these comparisons. The number of modules enriched in these GO terms is indicated on the Y axis in *log*2 scale. This suggests a conserved temporally coherent core immune response. Here, the temporal shift of the modules is shown with respect to Brevig and indicates a delay of these processes in Cal, Tx and NC responses with respect to the Brevig infection. Only GO terms with fewer than 100 annotated genes were considered. “Antigen processing and presentation of peptide antigen via MHC class I” was also a conserved GO term, but was enriched in very few modules and is omitted from the figure.

### 3.6 Identification of comparative differential subnetworks

Comparative differential subnetworks are a group of highly functionally related genes that show differences in their pattern of regulation in response to two perturbations being compared. Because their identification is a less constrained problem than conserved subnetwork discovery, the reliable selection of comparative differential modules is challenging. When identifying conserved subnetworks, the effects of noise in the data are mitigated by the requirement that common regulatory changes must be observed in different experiments. Methods that rely on pairwise gene interactions (41, 42) to reconstruct differential modules are limited by the fact that differential modules, by definition, allow inconsistencies across experiments. DYNAMO addresses this limitation by exploiting the functional modularity inherent in biological networks and leveraging the temporal dimension of time-course expression data in its time-lag optimization search. These aspects of the algorithm allow to better constrain the problem of differentially activated gene identification and improve selection of high confidence subnetworks.

We applied our method for differential module discovery to all pairs of influenza strain responses. Shown in Supplementary Figure 2 are the results of DYNAMO’s application to the Cal/NC comparison. DYNAMO identified many differential comparative modules at a wide range of subnetwork score values. Importantly, the curves tracking subnetwork discovery for the real and randomized data show substantial separation, with false discovery rate of *<* 10% for a broad range of subnetwork score cutoffs. This finding implies that the differential subnetworks discovered by DYNAMO had a low proportion of false positives and likely represent biologically meaningful differences in the responses to the Cal and NC influenza infections. As such, DYNAMO is able to overcome some of the challenges in differential subnetwork discovery and improve the reliability of differential subnetwork identification.

Overall, we found many fewer differential modules as compared to conserved identified among pairs of responses (Supplementary Table 1). For example, we found only 20 differential subnetworks for the pair of responses Brevig/Cal that shows 207 conserved subnetworks. Also, in contrast to the results for the conserved subnetwork discovery, the differential modules generally did not exhibit a dominant time lag. Although DYNAMO benefits from its ability to optimize over a set of time lags to capture divergent expression trends and thereby identifies many more differential modules, these time lags do not exhibit a singular consistency. These results suggest that differentially active modules do not show the temporal coherence that was observed with conserved subnetworks.

In the four comparisons of differential modules for a pandemic and a seasonal strain, a considerable overlap in the GO terms assigned to these subnetworks was observed. We found 31 processes enriched (*p <* 0.0001) among the subnetworks and annotated by GO terms that are shared across pairs. The GO enrichment analysis of differential modules for the Cal/NC comparison is shown in Figure 5. Although the cytokine mediated signaling pathway was implicated as enriched by both conserved and differential modules (see also Figure 4), the genes contributing to this annotation show little overlap. For example, contributing to the annotation in the conserved modules of the Brevig/Cal comparison are classical antiviral program genes, including the MX, OAS, and IFIT family genes (43, 44, 45). Enrichment of the cytokine signaling pathway in the differential modules is driven by immunomodulatory genes, such as IL6 (46). These results reveal that despite sharing GO annotations, conserved and differential modules have different compositions.

**Figure 5.**
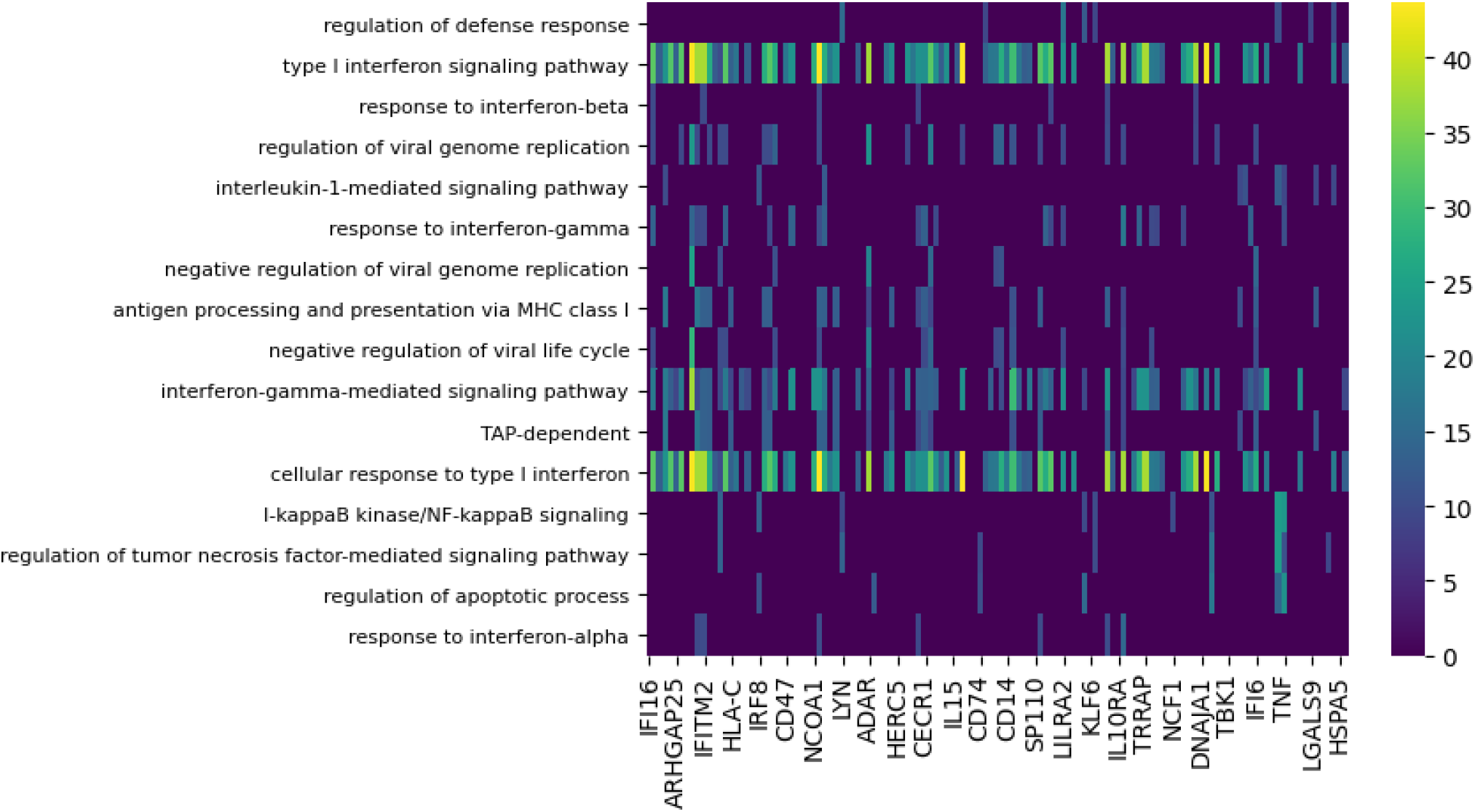
Functional enrichment analysis of differential subnetworks for the Cal/NC comparison using GO term enrichment with p-value cutoff 0.0001. The rows represent the GO terms and the columns are the individual subnetworks, indexed by their seed genes. The color scale indicates the level of significance of the GO term enrichment and is based on -log(pvalue). For clarity, the majority of GO terms that assign to fewer than seven subnetworks or that annotate more that 100 genes in GO are not shown. Notably, the same group of biological processes is enriched in differential subnetworks for the Brevig/Tx and Cal/Tx comparisons (data not shown).

Notably, the GO terms for antigen processing and apoptosis were enriched among the differential modules. Supplementary Figure 3 shows the genes implicated by the apoptosis-annotated subnetworks found to be differential between the seasonal and pandemic influenza infections. DYNAMO’s identification of apoptosis, known to be induced by influenza viruses (47, 27), as a differential process is consonant with other studies that show differences in global patterns of RNA degradation in response to seasonal and pandemic influenza infections (27).

### 3.7 DC antigen presentation differences after seasonal and pandemic influenza infection

The dendritic cell is a professional antigen presenting cell (48), raising the possibility that the seasonal and pandemic viruses differ in their modulation of this response. Dendritic cells, important mediators of innate and adaptive immunity, act by presenting antigens to T cells to initiate adaptive immune responses (48). Antigen presentation occurs either via direct presentation of digested viral peptides on the surface of infected cells or via cross-presentation of exogenous antigens by uninfected cells. Other studies (49) have found that influenza infection reduces both the efficiency of influenza antigen presentation and the ability of dendritic cells to cross-present antigens from other pathogens, such as bacteria, that cause co-infection during the course of influenza infection. In view of the centrality of antigen presentation by dendritic cells in the immunological response to influenza, the differences in antigen presentation between seasonal and pandemic influenza viruses inferred by the DYNAMO analysis might contribute to differences in the immunological and clinical response to these viruses.

To validate the hypothesis of the DYNAMO algorithm, we experimentally tested if infection with the two seasonal and one pandemic IAV strain resulted in differences in antigen presentation. Antigen presentation by professional APCs, such as dendritic cells, occurs via three signals (50). In T cell-DC interaction, Signal 1 is the interaction of an MHC-I or MHC-II molecule loaded with a processed part of a pathogen, with the T cell receptor of an antigen specific T cell. Signal 2 consists of a set of co-stimulatory markers (e.g CD86 on the DCs, which interacts with CD28 on T cells). Signal 2 is essential for T cell activation as presentation of a pathogen through Signal 1 alone leads to anergy of the specific T cells. Signal 3 consists of secreted factors, which influence the direction of the target T cells (eg Th1, Th2 or Th17). Here we used flow cytometry to quantify the induction of Signal 1 (MHC-II) and Signal 2 (CD86) after infection with one pandemic (Cal) and two seasonal (Tx and NC) H1N1 IAV strains. Consonant with the prediction of the DYNAMO algorithm, infection with the seasonal IAV strains resulted in lower expression of HLADR (Signal 1) and CD86 (Signal 2) in comparison with infection with the pandemic strain (Figure 6). (*p <* 0.05 for Cal/NC and *p <* 0.005 for Cal/Tx, using Student’s t-test).

**Figure 6.**
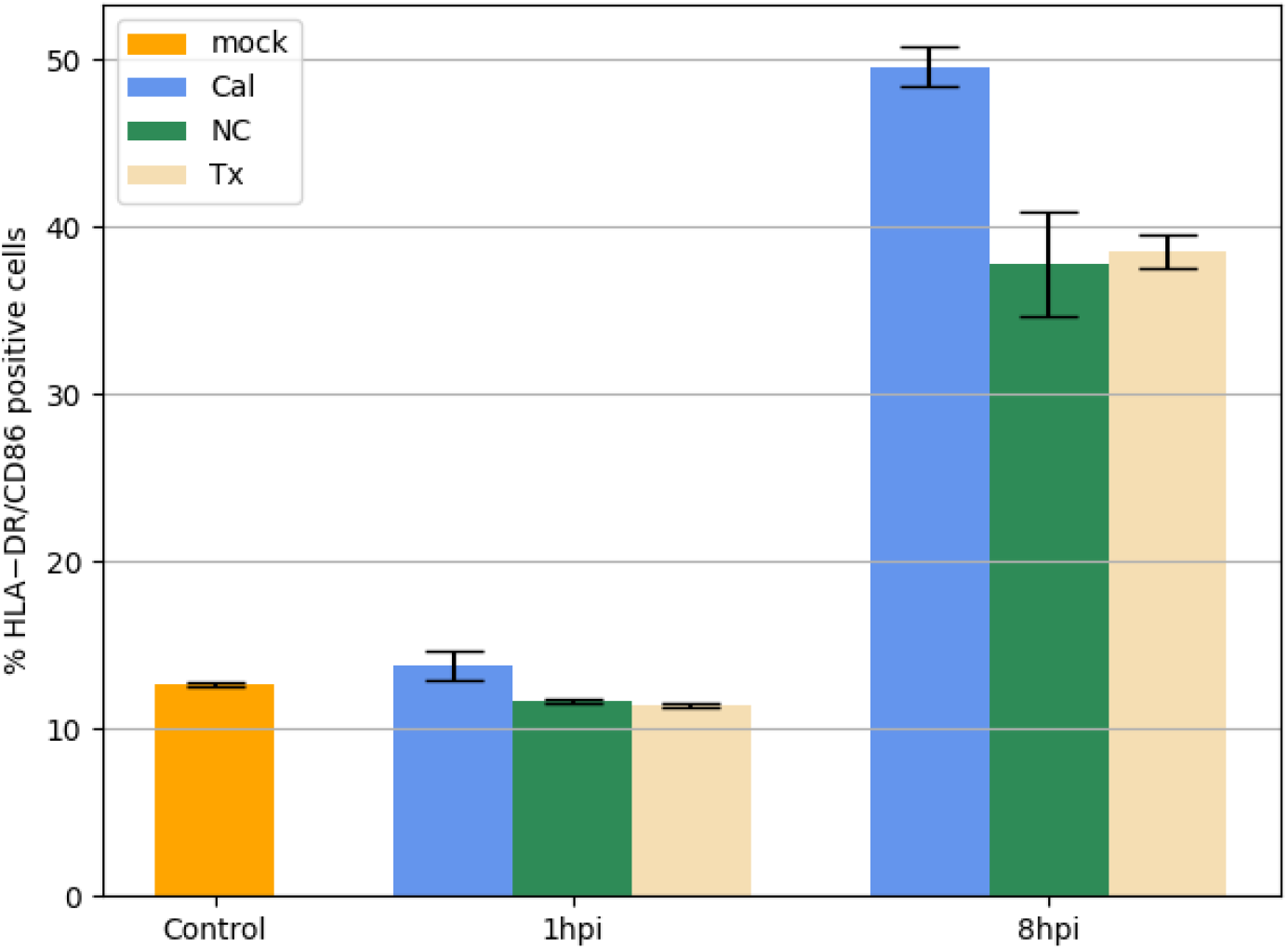
DC HLADR/CD86 surface expression following IAV infection. Percent of dendritic cells showing both HLADR and CD86 surface marker expression is plotted after infection with seasonal or pandemic IAV strains as measured by flow cytometry. All experiments were done in triplicates. Values shown are *mean± s*.*e*.*m*. The levels of surface marker expression was identical following mock and each virus infection at 1 h, and diverged at 8 h (*p <* 0.05, Student’s t-test), with higher surface marker expression following the pandemic Cal09 infection.

Overall, the application of DYNAMO to the seasonal and pandemic H1N1 influenza infection datasets derived insight into commonalities and differences in the regulation of functional modules and potential mechanisms of immune response modulation by the individual influenza virus strains.

## 4 DISCUSSION

In this study, we applied DYNAMO, a technique for discovery of comparative modules with different temporal dynamics or patterns of activation, to investigate host responses to infection by four different influenza virus strains and gain insight into the temporal and functional similarities and differences between them. We showed that the ability to search over multiple temporal lags allowed us to discover conserved temporally shifted mechanisms between different immune responses. Overall, we found remarkable temporally coherent conservation of a core group of immune processes that are crucial to infection control, such as cytokine signaling and specifically interferon signaling, in responses to all four viruses.

Our search for differential modules pointed to potential mechanistic differences among the seasonal and pandemic strains, discovering subnetworks that suggest a key role for apoptosis, a finding consonant with previous experimental work implicating apoptosis in the host response to influenza (47). Moreover, presence of differential subnetworks functionally annotated with antigen processing and presentation suggests an interesting potential direction for future experimental work.

Methodologically, the development of DYNAMO represents an important advance, which adds the element of temporal dynamics to the broad systems biology problem of functional subnetwork discovery (18). Our method builds upon the successes in the development of sophisticated integrative approaches that combine heterogeneous data to elucidate the modular functional architecture of the cell. DYNAMO is the first method to successfully exploit the temporal dimension of gene expression data for comparative module discovery and analysis.

While our algorithm development and successful application to the study of the immune response to multiple strains of the influenza virus is encouraging, a number of promising directions for further improvement of the method remain. The current version is restricted to expression data that is identically sampled and aligned. Since few datasets in the public domain share the same experimental design, relaxing this restriction, possibly using the time-warping algorithm (26), would make our method more broadly applicable. Furthermore, the approach is readily extended to simultaneously compare more than two datasets. Together, these improvements would enable the study of conserved and differential components of the response to infection by multiple pathogens, providing insight into the functioning of the host immune system and common and unique aspects of virus-host interactions, as well as facilitate comparative study of the pattern and timing of module activation elicited by other biological stimuli.

## AUTHOR CONTRIBUTIONS

Irina Nudelman and Daniil Kudrin developed and implemented the algorithm and conducted all data analysis. German Nudelman provided support for the development of the website. Raamesh Deshpande assisted with algorithm implementation and software development. Boris M. Hartmann conducted the experimental work. Steven H. Kleinstein, Chad L. Myers and Stuart C. Sealfon supervised the analyses and edited the manuscript. Elena Zaslavsky conceptualized and managed the research project, supervised the algorithm development and data analyses and wrote the manuscript.

## REFERENCES

1 Watanabe T, Kawaoka Y. Pathogenesis of the 1918 pandemic influenza virus. PLoS Pathog 7 (2011) e1001218.

2 Michaelis M, Doerr HW, Cinatl J Jindrich. Novel swine-origin influenza a virus in humans: another pandemic knocking at the door. Med. Microbiology and Immunology 198 (2009) 175–183.

3 Korth MJ, Tchitchek N, Benecke AG, Katze MG. Systems approaches to influenza-virus host interactions and the pathogenesis of highly virulent and pandemic viruses. Seminars in Immunology 25 (2013) 228–239.

4 Geiss GK, Salvatore M, Tumpey TM, Carter VS, Wang X, Basler CF, et al. Cellular transcriptional profiling in influenza A virus-infected lung epithelial cells: The role of the nonstructural NS1 protein in the evasion of the host innate defense and its potential contribution to pandemic influenza. PNAS 99 (2002) 10736–10741.

5 Baskin CR, García-Sastre A, Tumpey TM, Bielefeldt-Ohmann H, Carter VS, Nistal-Villán E, et al. Integration of clinical data, pathology, and cDNA microarrays in influenza virus-infected pigtailed macaques (macaca nemestrina). J. Vir. 78 (2004) 10420–10432.

6 Kash J, Tumpey T, Proll S, Carter V, Perwitasari O, Thomas M, et al. Genomic analysis of increased host immune and cell death responses induced by 1918 influenza virus. Nature 443 (2006) 578–581.

7 Kobasa D, Jones SM, Shinya K, Kash JC, Copps J, Ebihara H, et al. Aberrant innate immune response in lethal infection of macaques with the 1918 influenza virus. Nature 445 (2007) 319–323.

8 Lee SMY, Gardy JL, Cheung CY, Cheung TKW, Hui KPY, Ip NY, et al. Systems-level comparison of host-responses elicited by avian H5N1 and seasonal H1N1 influenza viruses in primary human macrophages. PLoS ONE 4 (2009) e8072.

9 Josset L, Belser JA, Pantin-Jackwood MJ, Chang JH, Chang ST, Belisle SE, et al. Implication of inflammatory macrophages, nuclear receptors, and interferon regulatory factors in increased virulence of pandemic 2009 H1N1 influenza A virus after host adaptation. J. Vir. 86 (2012) 7192–7206.

10 Ideker T, Ozier O, Schwikowski B, Siegel AF. Discovering regulatory and signalling circuits in molecular interaction networks. Bioinformatics 18 (2002) S233–S240.

11 Reiss D, Baliga N, Bonneau R. Integrated biclustering of heterogeneous genome-wide datasets for the inference of global regulatory networks. BMC Bioinformatics 7 (2006) 280.

12 Lan A, Smoly IY, Rapaport G, Lindquist S, Fraenkel E, Yeger-Lotem E. ResponseNet: revealing signaling and regulatory networks linking genetic and transcriptomic screening data. Nucleic Acids Research 39 (2011) W424–W429.

13 Leiserson MD, Blokh D, Sharan R, Raphael BJ. Simultaneous identification of multiple driver pathways in cancer. PLoS Comput Biol 9 (2013) e1003054.

14 Deshpande R, Sharma S, Verfaillie CM, Hu WS, Myers CL. A scalable approach for discovering conserved active subnetworks across species. PLoS Comput Biol 6 (2010) e1001028.

15 Waltman P, Kacmarczyk T, Bate A, Kearns D, Reiss D, Eichenberger P, et al. Multi-species integrative biclustering. Genome Biology 11 (2010) R96.

16 Gill R, Datta S, Datta S. A statistical framework for differential network analysis from microarray data. BMC Bioinformatics 11 (2010) 95.

17 Zhang B, Tian Y, Jin L, Li H, Shih IM, Madhavan S, et al. DDN: a caBIG analytical tool for differential network analysis. Bioinformatics 27 (2011) 1036–1038.

18 Mitra K, Carvunis AR, Ramesh SK, Ideker T. Integrative approaches for finding modular structure in biological networks. Nat Rev Genet 14 (2013) 719–732.

19 Luscombe N, Babu M, Yu H, Snyder M, Teichmann S, Gerstein M. Genomic analysis of regulatory network dynamics reveals large topological changes. Nature 431 (2004) 308?–312.

20 Przytycka TM, Singh M, Slonim DK. Toward the dynamic interactome: it’s about time. Briefings in Bioinformatics 11 (2010) 15–29.

21 Bar-Joseph Z, Gitter A, Simon I. Studying and modelling dynamic biological processes using time-series gene expression data. Nat Rev Genet 13 (2012) 552–564.

22 Gao S, Wang X. Identification of highly synchronized subnetworks from gene expression data. BMC Bioinformatics 14 (2013) s5.

23 Jin R, McCallen S, Liu C, Xiang Y, Almaas E, Xj Z. Identifying dynamic network modules with temporal and spatial constraints. Pac Symp Biocomput (2009), 203–214.

24 ElBakry O, Ahmad M, Swamy M. Inference of gene regulatory networks from time-series microarray data. NEWCAS Conference (NEWCAS), 2010 8th IEEE International (2010), 141–144.

25 Gao S, Hartman IV J, Carter J, Hessner M, Wang X. Global analysis of phase locking in gene expression during cell cycle: the potential in network modeling. BMC Systems Biology 4 (2010) 167.

26 Aach J, Church GM. Aligning gene expression time series with time warping algorithms. Bioinformatics 17 (2001) 495–508.

27 Hartmann BM, Thakar J, Albrecht RA, Avey S, Zaslavsky E, Marjanovic N, et al. Human dendritic cell response signatures distinguish 1918, pandemic and seasonal H1N1 influenza viruses. J Virol. 89 (2015) 10190–10205.

28 Benjamini Y, Hochberg Y. Controlling the false discovery rate: A practical and powerful approach to multiple testing. J. Royal Statistical Society 57 (1995) 289–300.

29 Huttenhower C, Haley EM, Hibbs MA, Dumeaux V, Barrett DR, Coller HA, et al. Exploring the human genome with functional maps. Genome Res. 19 (2009) 1093–1106.

30 Kanehisa M, Goto S, Kawashima S, Okuno Y, Hattori M. The KEGG resource for deciphering the genome. Nuc. Acids Res. 32 (2004) D277–D280.

31 Gorenshteyn D, Zaslavsky E, Fridbourg M, Park C, Wong AK, Tadych A, et al. Interactive big data resource to elucidate human immune pathways and diseases. Immunity 43 (2015) 605—-614.

32 Guan Y, Myers CL, Lu R, Lemischka IR, Bult CJ, Troyanskaya OG. A genomewide functional network for the laboratory mouse. PLoS Comput Biol 4 (2008) e1000165.

33 Myers CL, Robson D, Wible A, Hibbs MA, Chiriac C, Theesfeld CL, et al. Discovery of biological networks from diverse functional genomic data. Genome Biology 6 (2005) R114.

34 Zinman GE, Naiman S, O’Dee DM, Kumar N, Nau GJ, Cohen HY, et al. Moduleblast: identifying activated sub-networks within and across species. Nucleic Acids Research 43 (2015) e20.

35 Pietro Zoppoli MC Sandro Morganella. Timedelay-aracne: Reverse engineering of gene networks from time-course data by an information theoretic approach. BMC Bioinformatics 11 (2010).

36 Ashburner M, Ball CA, Blake JA, Botstein D, Butler H, Cherry JM, et al. Gene Ontology: tool for the unification of biology. Nat. Gen. 25 (2000) 25–29.

37 et al CE. Enrichr: interactive and collaborative html5 gene list enrichment analysis tool. BMC Bioinformatics 128 (2013).

38 Garcia-Sastre A, Durbin RK, Zheng H, Palese P, Gertner R, Levy DE, et al. The role of interferon in influenza virus tissue tropism. J. Vir. 72 (1998) 8550–8558.

39 Haller O, Kochs G, Weber F. The interferon response circuit: Induction and suppression by pathogenic viruses. Virology 344 (2006) 119 – 130.

40 Cillúniz C, Shinya K, Peng X, Korth MJ, Proll SC, Aicher LD, et al. Lethal influenza virus infection in macaques is associated with early dysregulation of inflammatory related genes. PLoS Pathog 5 (2009) e1000604.

41 Eddy JA, Hood L, Price ND, Geman D. Identifying tightly regulated and variably expressed networks by differential rank conservation (DIRAC). PLoS Comput Biol 6 (2010) e1000792.

42 Valcàrcel B, Würtz P, Seich al Basatena NK, Tukiainen T, Kangas AJ, Soininen P, et al. A differential network approach to exploring differences between biological states: An application to prediabetes. PLoS ONE 6 (2011) e24702.

43 Pitossi F, Blank A, Schröder A, Schwarz A, Hüssi P, Schwemmle M, et al. A functional GTP-binding motif is necessary for antiviral activity of Mx proteins. Journal of Virology 67 (1993) 6726–6732.

44 Melchjorsen F, Kristiansen H, Christiansen R, Rintahaka J, Matikainen S, Paludan S, et al. Differential regulation of the OASL and OAS1 genes in response to viral infections. Journal of Interferon & Cytokine Research 29 (2009) 199–208.

45 Diamond MS, Farzan M. The broad-spectrum antiviral functions of IFIT and IFITM proteins. Nat Rev Immunol 13 (2013) 46–57.

46 Dienz O, Rud J, Eaton S, Lanthier P, Burg E, Drew A, et al. Essential role of IL-6 in protection against H1N1 influenza virus by promoting neutrophil survival in the lung. Mucosal Immunol 5 (2012) 258–266.

47 Schultz-Cherry S, Krug RM, Hinshaw VS. Induction of apoptosis by influenza virus. Seminars in Virology 8 (1998) 491 – 495.

48 Guermonprez P, Valladeau J, Zitvogel L, Théry C, Amigorena S. Antigen presentation and T cell stimulation by dendritic cells. Ann. Rev. Imm. 20 (2002) 621–667.

49 Smed-Sorensen A, Chalouni C, Chatterjee B, Cohn L, Blattmann P, Nakamura N, et al. Influenza A virus infection of human primary dendritic cells impairs their ability to cross-present antigen to CD8 T cells. PLoS Pathog 8 (2012) e1002572.

50 Hufford MM, Kim TS, Sun J, Braciale TJ. The effector t cell response to influenza infection. Current topics in microbiology and immunology 386 (2015) 423–455.

51 Pavlovic J, Haller O, Staeheli P. Human and mouse Mx proteins inhibit different steps of the influenza virus multiplication cycle. J. Vir. 66 (1992) 2564–2569.

