## Supplemental Tables and Figures for "Comparing host module activation patterns and temporal dynamics in infection by influenza H1N1 viruses"

### Supplementary Materials

DYNAMO is an integrative method for comparative functional module discovery that uses time series gene expression data and a functional network to extract high confidence active subnetworks that exhibit either conserved or differential regulation between a pair of conditions. In this manuscript we developed the method and applied it to compare host immune responses to four different influenza A/H1N1 strain infections.

#### Subnetwork score calculation for conserved and differential search.

The derivation of the subnetwork scores used by the DYNAMO algorithm differs slightly for discovery of conserved and differential subnetworks. As can be seen in the top panel of Supplementary Figure 1, the distribution of correlations between expression profiles of a gene in two responses is shifted toward high positive values. Applying Fisher's Z-transform translates the correlations distribution to be approximately normal (middle panel). DYNAMO uses the transformed score in the conserved module search, setting the threshold at 1.5. For differential subnetworks discovery, the goal is to identify groups of genes that show low average correlations between the responses. To identify sets of genes with low correlations from different distributions, we standardize the Fisher's Z-transformed values of each distribution, setting its mean to zero, and select genes with values that are some number of standard deviations below the mean. We use -1.5 as the subnetwork score cutoff in the differential module search, which seeks to minimize the subnetwork score. Equivalently, we run a maximization procedure, having multiplied the scores by -1, and using 1.5 as the threshold. Using these settings allows for many modules to be discovered. Low confidence modules, as determined by the application of the randomization procedure and application of the confidence threshold, are removed.

#### Discovery of high confidence differential subnetworks.

We applied our method for differential comparative module discovery to all pairs of influenza strain responses. The results of our comparisons are listed in Supplementary Table 1. Shown below in Supplementary Figure 2 are the results of DYNAMO's application to the Cal/NC comparison for differential module discovery.

Furthermore, in the four comparisons of differential modules for a pandemic and a seasonal strain, a considerable overlap in the GO terms assigned to these subnetworks was observed. We highlight the presence of apoptosis among these terms as a differential process that is consonant with other studies and underlies differences in global patterns of RNA degradation in response to seasonal and pandemic influenza infections. Supplementary Figure 3 shows the genes implicated by the apoptosis-annotated subnetworks found to be differential between the seasonal and pandemic influenza infections.

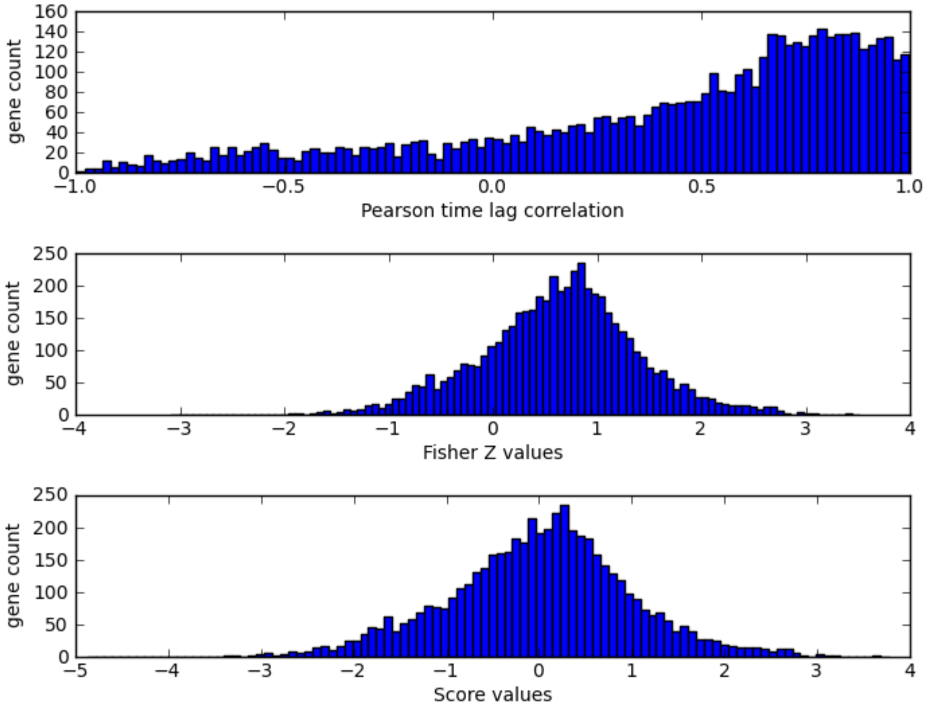


Supplementary Figure 1: Distributions of raw and transformed correlations for the Cal/Brevig pair at the 80 minute time lag. The top panel shows the raw correlations for each gene between its time courses in the two infections. Applying the Fisher's Z-transform gives the distribution a bell shape and shows its mean at approximately 0.9. These transformed correlations were used in the conserved search. The third panel shows the standardization of the distribution with the mean shifted to 0. These standardized scores were used in the differential module search.

Table 1: **Differential comparative subnetworks found in comparisons of responses to influenza strain infections:**

| pair | number of subnetworks | average size |
|---|---|---|
| Brevig/Cal | 20 | 18 |
| Cal/Tx | 90 | 23 |
| Brevig/Tx | 45 | 21 |
| Cal/NC | 127 | 23 |
| NC/Tx | 149 | 22 |
| Brevig/NC | 62 | 22 |

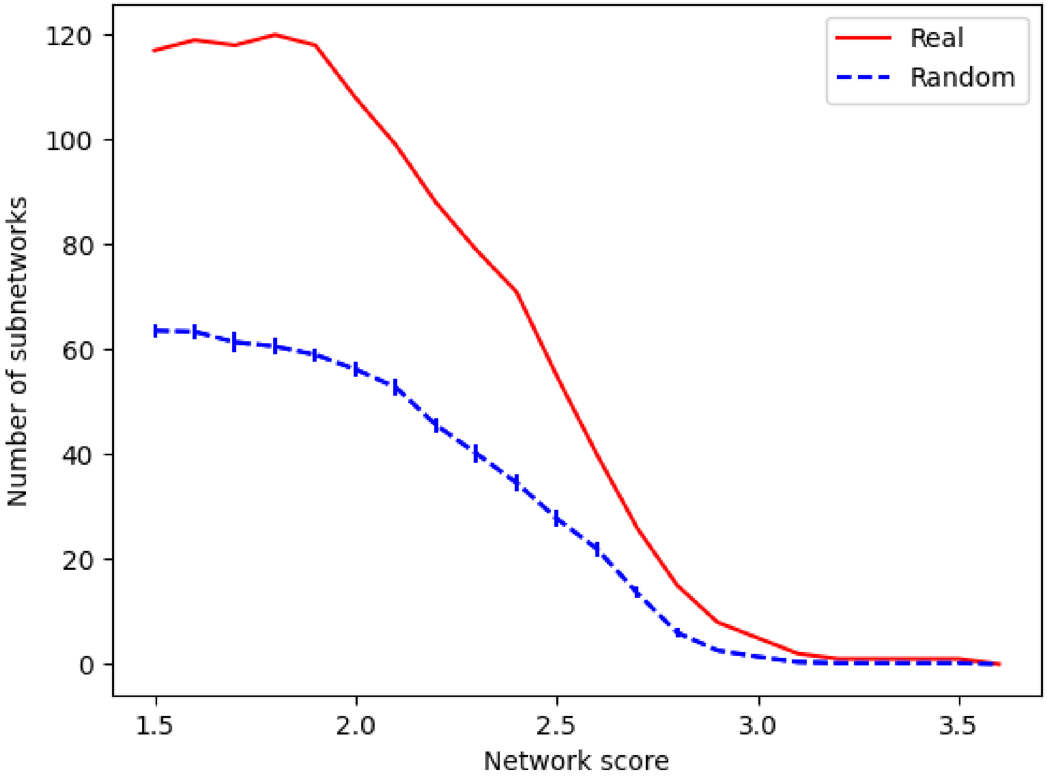


Supplementary Figure 2: Assessment of the false discovery rate in DYNAMO's differential comparative module search for the NC/Cal comparison. The solid line represents the number of differential subnetworks discovered across a wide range of subnetwork expression scores, while the dashed line shows results of the randomization analysis, averaged over five randomization instances. All discovered modules are reported, regardless of the confidence threshold.

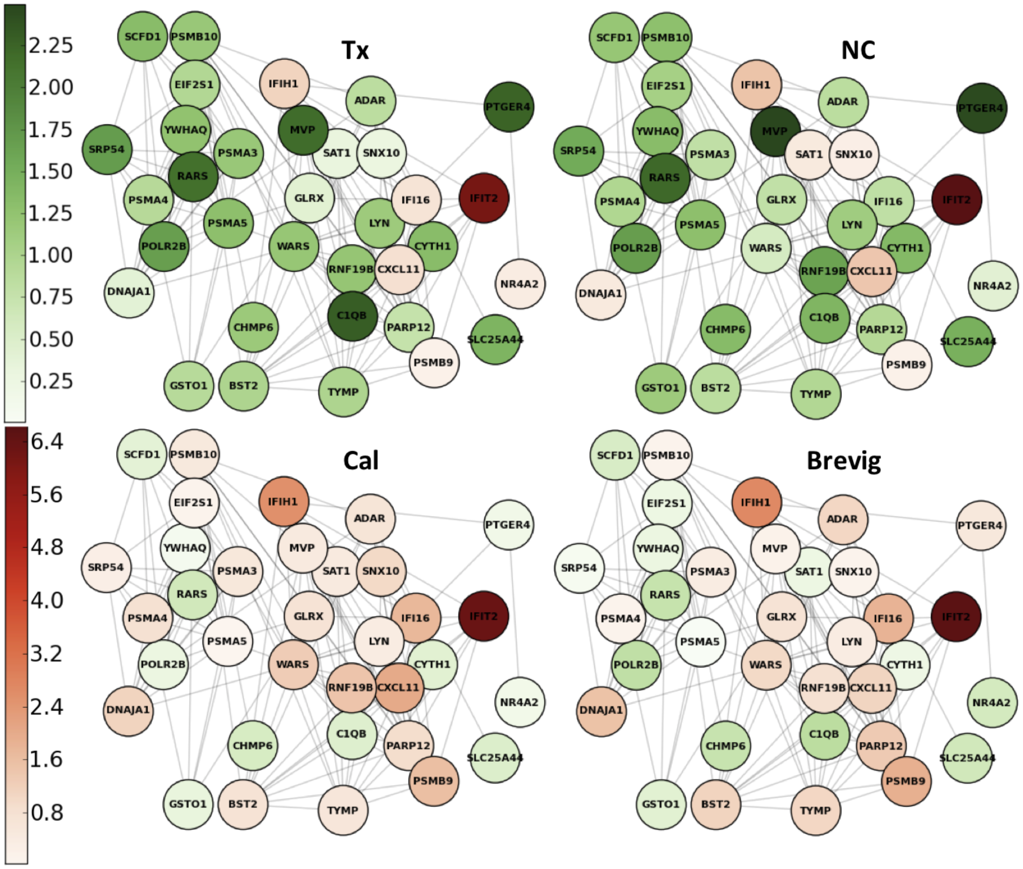


Supplementary Figure 3: A summary figure showing a subgraph with expression values of module genes that implicate apoptosis-related subnetworks in multiple comparisons between a pandemic and a seasonal strain infection response performed using DYNAMO. The edges between the seed genes come from the immune-specific functional network. Note the differential expression pattern of the apoptosis subnetwork seed genes observed following infection by pandemic and seasonal viruses. Each node is colored based on its maximal fold-change along the time course in either positive or negative direction. Red and green indicate positive and negative absolute maximal fold changes respectively.
